# Genetic variation for root architectural traits in response to phosphorus deficiency in mungbean at the seedling stage

**DOI:** 10.1101/719773

**Authors:** Venkata Ravi Prakash Reddy, M. Aski, G.P. Mishra, H.K. Dikshit, Akanksha Singh, Renu Pandey, Madan Pal, Gayacharan, Vinita Ramtekey, Priti, Neha Rai

## Abstract

Roots enable the plant to survive in natural environment by providing anchorage and acquisition of water and nutrients. In this study, 153 mungbean genotypes were studied to compare root architectural traits under normal and low phosphorus conditions. Significant variations, medium to high heritability, near normal distribution and significant correlations were observed for studied root traits. Total root length (TRL) was positively correlated with total surface area (TSA), total root volume (TRV), total root tips (TRT) and root forks. The first two principal components explained the 79.19 % and 78.84% of the total variation under normal and low phosphorus conditions. TRL, TSA and TRV were major contributors of variation and can be utilized for screening of phosphorus uptake efficiency at seedling stage. Released Indian mungbean varieties were found to be superior for root traits than other genotypic groups. Based on comprehensive phosphorus efficiency measurement, IPM-288, TM 96-25, TM 96-2, M 1477, PUSA 1342 were found to be best five highly efficient genotypes whereas M 1131, PS-16, Pusa Vishal, M 831, IC 325828 were highly inefficient genotypes. These identified highly efficient lines are valuable genetic resources for phosphorus uptake efficiency that could be used in mungbean breeding programme.

## Introduction

Mungbean is important warm season grain legume grown in more than 6 million hectare area [1] for its protein rich seeds. Mungbean seeds are rich source of iron [2] and Vitamin C and folates [3]. Cultivation of this crop improves soil fertility through biological nitrogen fixation [4]. Mungbean is cultivated on marginal lands resulting in poor growth, development and yield. Fertilizer management is important for realizing the potential yield in marginal lands of the crop [5]. Nitrogen and phosphorus are the important macronutrients required for the crop. In mungbean 80-90% nitrogen requirement is met through biological N_2_ fixation mechanism [6]. Mungbean requires 48.1 kg P_2_O_5_ for producing one ton of grains [7]. Under tropical and subtropical conditions, phosphorus is the main yield limiting factor [8].

Globally, by the year 2020, phosphorus fertilizer requirement is expected to reach 64.68 million tonnes, whereas, the estimated supply is 53.08 million tonnes and demand for phosphorus fertilizer requirement is increasing annually by 2.2% on average from 2015-2020 [9]. US, China and Morocco are the leading producers of phosphate fertilizer [10]. Expecting future domestic demands US and China have stopped the export of rock phosphate to other countries [11]. Phosphorus deficiency leads to higher root/shoot ratio as shoot growth is relatively more affected in comparison to root growth. It also causes stunted growth and foliage turns dark green colour due to accumulation of starch and sugars in the leaves. Deficiency in leaves disturbs the photosynthetic machinery and electron transport chain through repression of orthophosphate concentration in chloroplast stroma inhibiting ATP synthase activity [12]. Phosphorus is key component of nucleic acids and plant hormones and determines the yield and quality of a crop [13, 14]. Phosphorus deficiency in soil can be overcome by phosphorus fertilizer application, excess application leads to delayed formation of reproductive organs [15]. Uptake of phosphorus from soil is complex as the phosphorus is bound to calcium in alkaline soils and iron and aluminium in acid soils [16].

Plants exposed to phosphorus deficiency can activate a range of mechanisms that either result in increased acquisition of phosphorus from soil or more efficient utilization of internal phosphorus [8]. Plant undergoes modification of root morphology, change of root physiology, increased expression of high affinity phosphorus transporters and increased root microbial association. Change in root architecture exploits the soil space and enhanced root-soil contact to increase phosphorus uptake [17-19]. Root is the indispensable organ of the plant for absorption of nutrients and water by expanding its surface area and enhancement of explored soil volume [20]. Alteration of root architecture in response to phosphorus deprivation mainly depends on localized P concentration, sensitivity to or transport of growth regulators such as auxins, ethylene, cytokinins, sugars, nitric oxide, reactive oxygen species and abscisic acid (ABA) [21].

Genetic variation in plant root architecture can be exploited to improve the nutrient and water use efficiency under difficult growing conditions [22]. Root surface area, volume, biomass and root caboxylate exudation capacity were reported to be significantly higher in phosphorus efficient mungbean genotype compared to inefficient genotype [23]. Significant contribution of root length, root volume, surface area and number of lateral root towards phosphorus uptake at 45 days after sowing was observed in blackgram [24]. In rice, root hair length and density significantly increased in all tested genotypes under low phosphorus conditions [25]. Shen *et al*. [26] stressed on maintaining of root biomass and root length to cope with deficiency of phosphorus in wheat.

Although phosphorus deficiency can affect the crop growth throughout the season, phenotypic evaluation at seedling stage is an attractive approach because of high throughput and low cost method that saves time and space [27]. Stress gradient hypothesis [28, 29] proposes that the fate of seedlings determines the structure and dynamics of plant population. Current digital image analysis enables accurate analysis of plant root system and is time and labour saving technology [30, 31]. Considering the role of root architecture in phosphorus uptake the present study was designed to (i) characterize the phenotypic variation for morphological root traits in 153 mungbean lines (ii) identify the root related traits accounting for most of the variation among the tested mungbean lines (iii) evaluate the efficiency of mungbean lines under normal and low phosphorus conditions.

## Methods and material

### Plant materials and plant growth conditions

One hundred and fifty three mungbean lines including 41 Indian released varieties (IRV), 44 Advanced Breeding lines (ABL) and 68 Germplasm lines (GL) were studied for root architecture characteristics under normal and low phosphorus conditions (**Supplemental Table 1)**. The experiment was conducted in a National Initiative on Climate Resilient Agriculture (NICRA)-controlled environment facility of the Indian Agricultural Research Institute, New Delhi, India from December, 2017 to September, 2018. In this chamber, the growth conditions were maintained as: 30/18 °C day/night temperature, photoperiod of 12 h with photon flux density of 450 µmol m^−2^ s^−1^ (PAR) and relative humidity at 90 %. For screening under hydroponics, mungbean seeds were surface sterilized with 0.1% (w/v) HgCl_2_ for 3 minutes and rinsed 3 times with double distilled water and wrapped in germination paper. Upon emergence of cotyledonary leaves after 5 days, seedlings of uniform size and without visible root injuries were transferred to modified Hoagland solution. Composition of basal nutrient solution was MgSO_4_ (1mM), K_2_SO_4_ (0.92 mM), CaCl_2_.2H_2_O (0.75 mM), KH_2_PO4 (0.25 mM), Fe-EDTA (0.04mM), Urea (5 mM), and micronutrients [H_3_BO_3_ (2.4μM), MnSO_4_ (0.9μM), ZnSO_4_ (0.6μM), CuSO_4_ (0.62μM), and Na_2_MoO_4_ (0.6μM)] [32]. Concentration of P was maintained with two P levels: normal P (250 μM) and low P (3μM). pH of the nutrient solution was maintained to 6.0 with 1 M KOH or 1 M HCl. Seedlings were supported on a 2” thick thermocol sheet with holes made at 5 × 5 cm plant-to-plant and row-to row distance. This sheet was fitted in to plastic containers (30 × 45 × 15 cm) with 10 L of basal nutrient solution. Forty five seedlings were raised in one such container and fifteen genotypes were screened at a time with three replicates for each genotype. The solution was aerated regularly by aquarium air pump and replaced on alternate days.

**Table 1:**
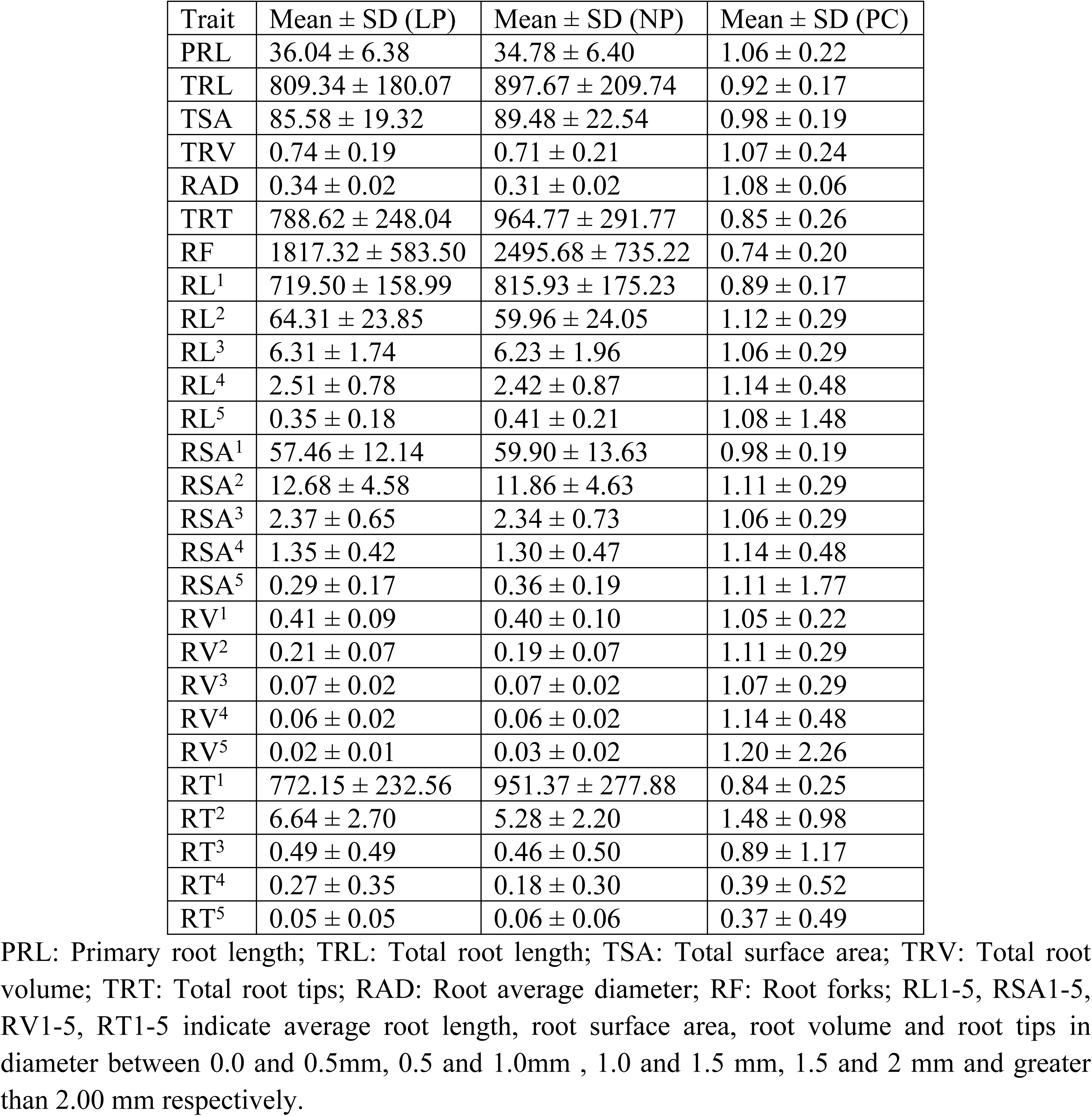
Mean value, standard deviation (SD) of traits investigated under two phosphorus regims and the phosphorus efficiency-coefficient (PC) of each trait

### Root measurements

The data for root traits and shoot traits was recorded on twenty one days old seedlings raised under low and sufficient phosphorus conditions. The complete root system was isolated from each plant and placed on a tray with no overlapping of any roots. Roots were scanned using root scanner (Epson professional scanner) and images were analyzed using WinRhizo Pro software (Regent Instrument Inc., Quebec, Canada). The following root parameters were measured: primary root length (PRL), total root length (TRL), total surface area (TSA), total root volume (TRV), root average diameter (RAD), total root tips (TRT), root forks (RF). WinRhizo also generated additional output that categorizes root traits [root length (RL), root surface area (RSA), root volume (RV) and no. of root tips (RT)] into five root diameter classes: 0-0.5 mm, 0.5-1.0 mm, 1.0-1.5 mm, 1.5-2.0 mm and >2.0 mm. Primary root length was measured manually using scale.

### Statistical analysis

The data was subjected to descriptive statistics including mean, standard deviation, coefficient of variation, analysis of variation, heritability and Pearson’s correlation were calculated for tested traits under normal and low phosphorus conditions using STAR (Statistical Tool for Agricultural Research) 2.1.0 software [33]. 153 mungbean lines were classified into three different categories based on their performance: (i) low performing lines with non-desirable root characteristics *(≤ X – SD*), (ii) medium performing lines *(≥X – SD*) to *(≤X* + *SD*), and (iii) high performing genotypes with desirable traits *(≥X* + *SD*) [34, 35]. A polymorphic diversity index, Shannon-Weaver diversity index (H’), was calculated for each trait [36, 37] using the formula

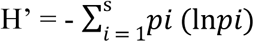

Where *pi* is the proportion of individuals belonging to the i^th^ species and s is the total number of species.

Principal component analysis (PCA) was performed to identify traits contributing most of variation in tested mungbean lines using STAR 2.1.0 software. A comprehensive phosphorus efficiency measurement value (P value) was used to estimate the efficiency capability of all tested mungbean lines. The P value was calculated across traits to evaluate mungbean phosphorus efficiency by using the formulas described below [38, 39].

The Phosphorus efficiency coefficient (PC) was calculated as the ratio of the data derived from the low phosphorus (LP) and normal phosphorus (NP) treatment of the same line for each trait using the following equation.

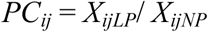

Where *PC*_*ij*_ is the phosphorus efficiency coefficient of the trait (*j*) for the cultivar (*i*); *X*_*ijLP*_ and *X*_*ijNP*_ are the value of the trait (*j*) for the cultivar (*i*) evaluated under low phosphorus (LP) and normal phosphorus (NP) treatments, respectively.

Fuzzy subordination method could be used to analyze the phosphorus efficiency completely and avoid the shortage of single index. The membership function of a fuzzy set is a generalization of the indicator function in classical sets; it represents the degree of truth as an extension of valuation [40]. *U*_*ij*_ stands for the membership function value of phosphorus efficiency (MFVP) that indicates a positive correlation between trait and phosphorus efficiency.

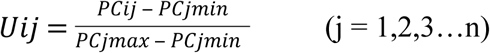

Where *U*_*ij*_ is the membership function value of the trait (*j*) for the cultivar (*i*) for phosphorus efficiency; *PC*_*jmax*_ is the maximum value of the phosphorus efficiency coefficient for the trait (*j*); *PC*_*jmin*_ is the minimum value of *PC*_*j*_.

Comprehensive phosphorus efficiency measurement was made using the formula:

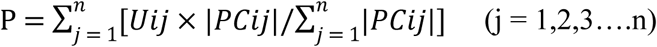

Where P is the comprehensive phosphorus efficiency measurement of each mungbean line under LP condition. Based P value all mungbean lines were classified into five groups, highly efficient, efficient, moderate efficient, inefficient and highly inefficient.

## Results

### Response of root traits to phosphorus stress

The study of 153 mungbean genotypes for root traits under normal phosphorus (NP) and low phosphorus (LP) conditions revealed high variation for means for the studied traits (**Table 1**). Compared to NP condition, the mean values of primary root length, total root volume and root average diameter were high in LP condition. For phosphorus efficiency coefficient (PC), root average diameter showed high mean value followed by total root volume and primary root length. The mean values of root length (RL), root surface area (RSA), root volume (RV) and no. of root tips (RT) in five root diameter classes: 0-0.5 mm, 0.5-1.0 mm, 1.0-1.5 mm, 1.5-2.0 mm and >2.0 mm exhibited variation under NP and LP conditions. RL and RSA revealed PC value above 1 for all root diameters except 0-0.5mm. Phosphorus efficiency coefficient (PC) was above 1 for RV at all root diameters indicating increase in root volume in LP condition. RT were higher under LP at root diameter of 0.5-1.0 mm.

### Genetic variation and broad sense heritability studies

ANOVA analysis revealed highly significant variation among the genotypes for seven traits (PRL, TRL, TSA, TRV, RAD, TRT and RF) evaluated under two phosphorus regimes (**Table 2**). The study revealed highly significant variation among the evaluated traits at two phosphorus conditions. The highly significant interaction between genotype and phosphorus treatment indicates that genotypes were significantly affected for studied root traits at different phosphorus regimes. The level of variation for studied seven phosphorus uptake efficiency traits is presented as **Fig. 1**. Histogram of frequency distribution revealed near normal distribution of root traits evaluated in the study. The coefficient of variation for seven investigated traits ranged from 4.64% (root average diameter) to 16.01% (root forks). The broad sense heritability for the studied traits ranged from 0.59 to 0.79. The highest broad sense heritability was observed in root average diameter (0.79) followed by total root volume (0.78) and lowest was observed in primary root length (0.59). The widely used indicator, root average diameter was highly heritable, suggesting that it is a reliable parameter for phosphorus efficiency.

**Table 2:** Analysis of variance for the tested traits under two phosphorus regimes.

| Variation source | df | Means of squares |  |  |  |  |  |  |
| --- | --- | --- | --- | --- | --- | --- | --- | --- |
|  |  | PRL | TRL | TSA | TRV | RAD | TRT | RF |
| Replication (R) | 2 | 39.48 | 83366.00 | 874.45 | 0.08 | 0 | 21779.66 | 364416.28 |
| Genotype (G) | 152 | 174.33*** | 183719.63*** | 2151.76*** | 0.20*** | 0.00*** | 321101.87*** | 2094494.45*** |
| Phosphorus (P) | 1 | 376.63*** | 1789765.97*** | 3503.19*** | 0.18** | 0.14*** | 7121141.19*** | 105608699.30*** |
| G × P | 152 | 70.89*** | 45566.49*** | 491.44*** | 0.04*** | 0.00*** | 118860.09*** | 548698.57*** |
| Error | 610 | 11.45 | 10418.26 | 117.79 | 0.01 | 0.00 | 18651.24 | 119218.02 |
| CV (%) |  | 9.55 | 11.96 | 12.40 | 15.26 | 4.64 | 15.58 | 16.01 |
| Heritability |  | 0.59 | 0.75 | 0.77 | 0.78 | 0.79 | 0.63 | 0.74 |

df: degree of freedom; PRL: Primary root length; TRL: Total root length; TSA: Total root surface area; TRV: Total root volume; RAD: Root average diameter; TRT: Total root tips; RF: Root forks; \*, \*\* and \*\*\* significant at $P < 0.05$ , $P < 0.01$ and $P < 0.001$ respectively.

**Fig.1.**
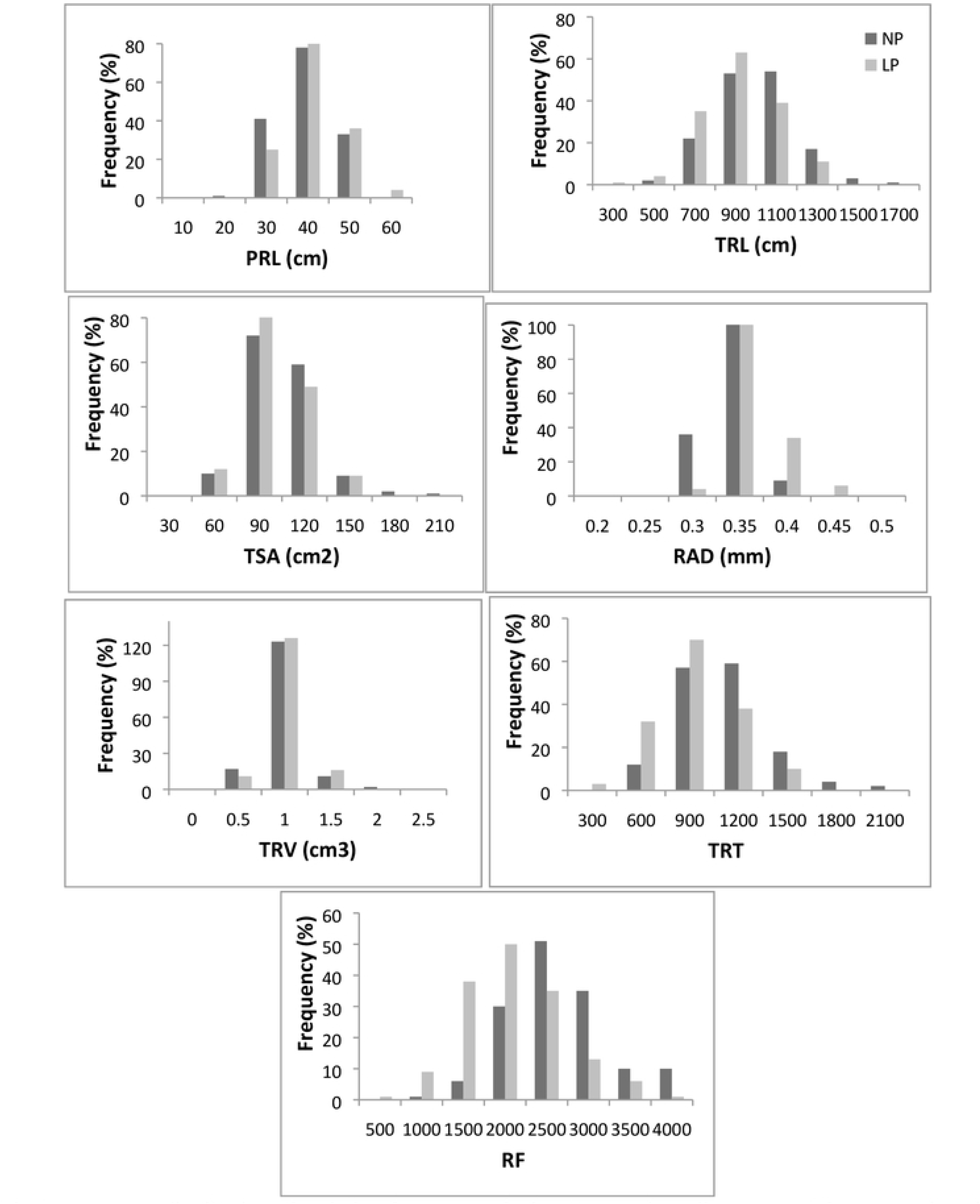
Frequency distribution of variation for seven traits in 153 mungbean lines. PRL: Primary root length; TRL: Total root length, TSA: Total root surface area; RAD: Root average diameter; TRV: Total root volume; TRT: Total root tips; RF: Root forks; NP: Normal phosphorus condition; LP: Low phosphorus condition.

### Genetic correlations among tested traits

Pearson correlations coefficients among all the traits under two phosphorus regimes were analyzed and significant correlations (p<.001) were observed between pairs of traits (**Table 3**). Under NP condition, highly significant and positive correlation was recorded between TRL and TSA (r = 0.953), TSA and TRV (r = 0.953) followed by TRL and TRV (r = 0.855). RF exhibited highly significant correlation with TRL, TSA and TRV. Whereas under LP condition, highly significant and positive correlation was observed for TRL and TSA (r = 0.951) followed by TSA and TRV (r = 0.929). Under both NP and LP conditions, TRV showed highly significant and positive correlation with all other tested traits. PRL and TRL showed significant and positive correlations with all other tested traits except root average diameter under both phosphorus regimes. Root average diameter showed significant negative correlation with total root tips and root forks under low phosphorus condition.

**Table 3:**
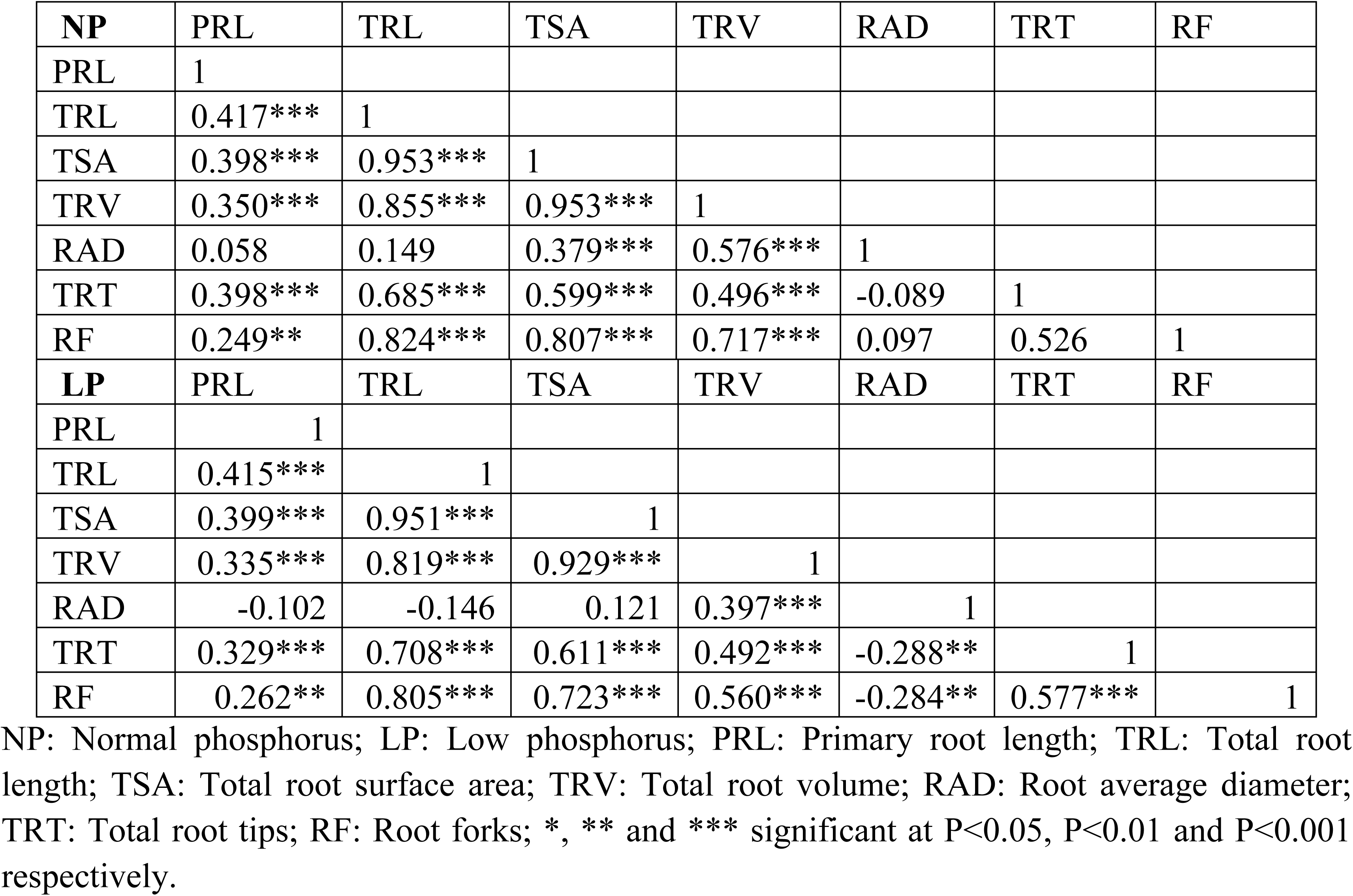
Genetic correlations among tested traits under normal and low phosphorus conditions.

### Comparison of root traits in different diameter classes under two phosphorus regimes

The proportion of roots in each diameter class was calculated as percentage of the total for each across different genotype groups under two phosphorus regimes (**Table 4**). The results revealed significantly higher percentage for studied root traits (RL, RSA, RV and RT) in 0-0.5 mm diameter class as compared to other diameter classes across different genotype groups under two phosphorus regimes (**Fig.2**). For root diameter classes 0.5-1.0 mm, 1.0-1.5 mm and 1.5-2.0 mm RL, RSA and RT percentage was higher in LP condition in comparison to NP condition. For diameter class of 0-0.5 mm and >2.0 mm, RL, RSA and RV recorded higher percentage in NP condition as compared to LP condition.

**Fig.2.**
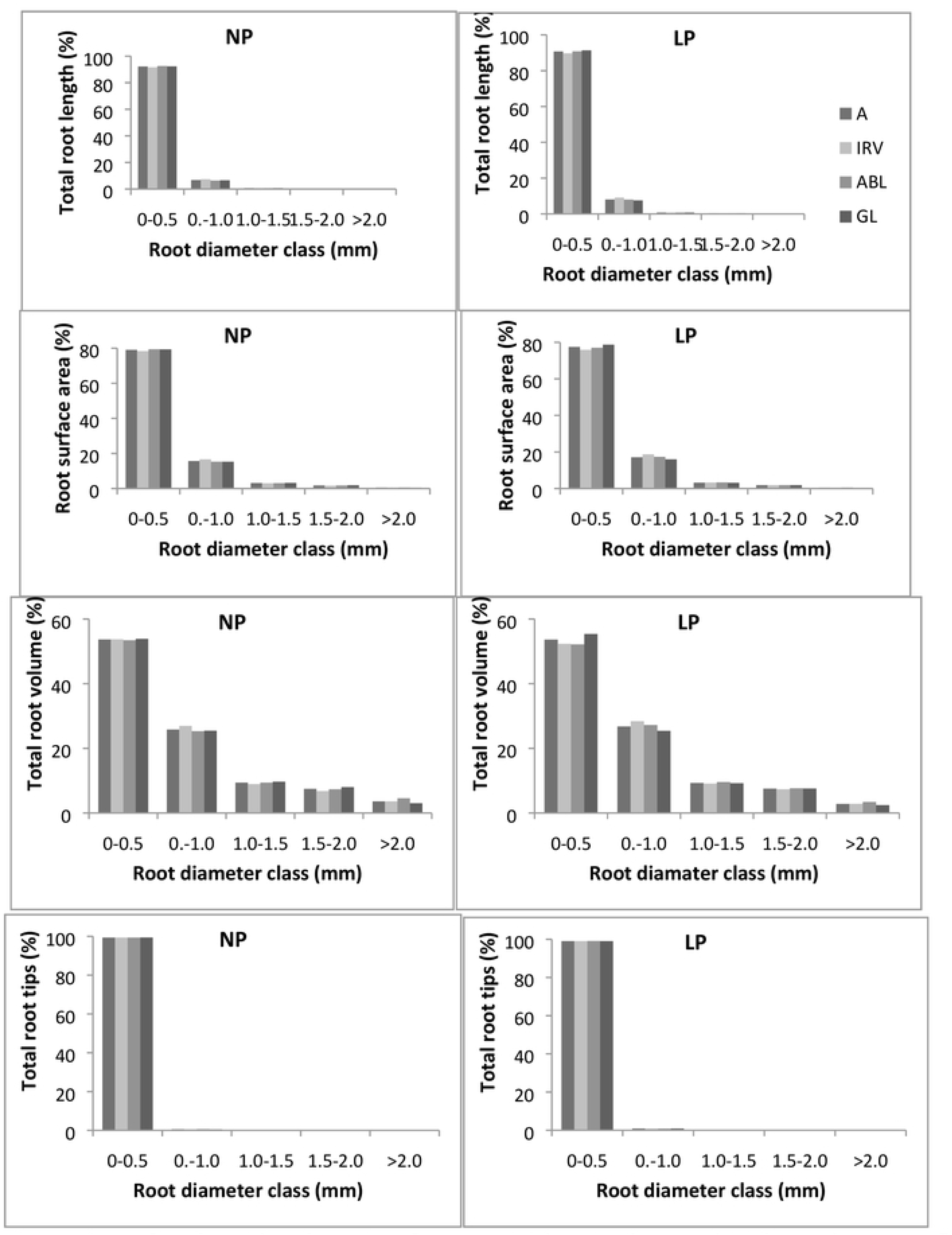
Percentage of total root length, root surface area, total root volume and total root tips across five diameter classes assessed under two phosphorus regimes.

**Table 4:** Percentage distribution of root traits across five root diameter classes under normal and low phosphorus conditions in different groups

| Traits | Treatment | Root diameter class (mm) |  |  |  |  |  |  |  |  |  |  |  |  |  |  |  |  |  |  |  |
| --- | --- | --- | --- | --- | --- | --- | --- | --- | --- | --- | --- | --- | --- | --- | --- | --- | --- | --- | --- | --- | --- |
|  |  | 0-0.5 |  |  |  | 0.5-1.0 |  |  |  | 1.0-1.5 |  |  |  | 1.5-2.0 |  |  |  | >2.0 |  |  |  |
|  |  | A | IRV | ABL | GL | A | IRV | ABL | GL | A | IRV | ABL | GL | A | IRV | ABL | GL | A | IRV | ABL | GL |
| RL (%) | NP | 92.20 | 91.61 | 92.58 | 92.32 | 6.78 | 7.39 | 6.43 | 6.62 | 0.70 | 0.69 | 0.68 | 0.73 | 0.27 | 0.25 | 0.26 | 0.29 | 0.05 | 0.05 | 0.05 | 0.04 |
|  | LP | 90.73 | 89.68 | 90.80 | 91.30 | 8.11 | 9.13 | 8.03 | 7.57 | 0.80 | 0.82 | 0.80 | 0.77 | 0.32 | 0.32 | 0.31 | 0.32 | 0.04 | 0.05 | 0.05 | 0.04 |
| RSA (%) | NP | 79.06 | 78.32 | 79.41 | 79.30 | 15.66 | 16.64 | 15.27 | 15.28 | 3.09 | 2.98 | 3.06 | 3.17 | 1.72 | 1.57 | 1.69 | 1.83 | 0.48 | 0.48 | 0.57 | 0.42 |
|  | LP | 77.48 | 75.85 | 77.04 | 78.73 | 17.10 | 18.70 | 17.34 | 16.00 | 3.20 | 3.22 | 3.30 | 3.12 | 1.82 | 1.82 | 1.84 | 1.81 | 0.40 | 0.42 | 0.48 | 0.35 |
| RV (%) | NP | 53.69 | 53.72 | 53.41 | 53.85 | 25.82 | 26.94 | 25.29 | 25.43 | 9.39 | 8.96 | 9.38 | 9.68 | 7.48 | 6.78 | 7.37 | 8.00 | 3.62 | 3.60 | 4.55 | 3.04 |
|  | LP | 53.63 | 52.32 | 52.15 | 55.37 | 26.74 | 28.40 | 27.21 | 25.42 | 9.29 | 9.09 | 9.59 | 9.22 | 7.53 | 7.37 | 7.63 | 7.56 | 2.82 | 2.83 | 3.41 | 2.43 |
| RT (%) | NP | 99.38 | 99.37 | 99.34 | 99.40 | 0.55 | 0.55 | 0.60 | 0.52 | 0.05 | 0.06 | 0.04 | 0.04 | 0.02 | 0.02 | 0.02 | 0.02 | 0.01 | 0.01 | 0.01 | 0.01 |
|  | LP | 99.04 | 99.02 | 99.08 | 99.04 | 0.85 | 0.86 | 0.83 | 0.86 | 0.06 | 0.08 | 0.05 | 0.06 | 0.03 | 0.04 | 0.03 | 0.03 | 0.01 | 0.01 | 0.01 | 0.01 |

NP: Normal phosphorus; LP: Low phosphorus; A: All 153 mungbean lines; IRV: Indian Released varieties; ABL: Advanced breeding lines; GL: Germplasm lines; RL: Root length; RSA: Root surface area; RV: Root volume; RT: Root tips;

### Diversity pattern with respect to different groups

A comparison of the root morphology of the different groups showed clear variation for all studied root traits. The mungbean lines in the IRV, ABL and GL groups were classified into three categories, namely low, medium and high performance (Table 5). For all studied traits higher number of genotypes were classified in medium group. Higher number of genotypes were grouped in high group for PRL, TRL, TRV, RAD and TRT under LP condition. Under LP condition, 8(20%), 7(16%) and 10 (15%) lines from IRV, ABL and GL groups showed larger TRL. For all traits except TSA, the GL group had a lower proportion of lines with high performance (≥X+SD) than either the IRV or ABL groups under two phosphorus regimes. Except for the trait TRL, ABL group had lower proportion of lines with high performance ((≥X+SD) than IRV group under two phosphorus regimes. This indicates that more lines with desirable root traits present in the order IRV>ABL>GL groups under the two phosphorus regimes.

**Table 5:** The Shannon-Weaver diversity index (H’) and performance categories under two different phosphorus conditions.

| Traits | Treatment | All 153 mungbean lines |  |  |  | Indian Released varieties (IRV) |  |  |  | Advanced breeding lines (ABL) |  |  |  | Germplasm lines (GL) |  |  |  |
| --- | --- | --- | --- | --- | --- | --- | --- | --- | --- | --- | --- | --- | --- | --- | --- | --- | --- |
|  |  | Low | Medium | High | H' | Low | Medium | High | H' | Low | Medium | High | H' | Low | Medium | High | H' |
| PRL | NP | 25 | 102 | 26 | 0.87 | 5 | 28 | 8 | 0.85 | 8 | 30 | 6 | 0.84 | 14 | 44 | 10 | 0.89 |
|  | LP | 22 | 103 | 28 | 0.83 | 4 | 29 | 8 | 0.79 | 5 | 29 | 10 | 0.86 | 11 | 48 | 9 | 0.81 |
| TRL | NP | 21 | 112 | 20 | 0.77 | 5 | 29 | 7 | 0.80 | 6 | 33 | 5 | 0.73 | 11 | 50 | 7 | 0.75 |
|  | LP | 23 | 106 | 24 | 0.86 | 5 | 28 | 8 | 0.84 | 4 | 33 | 7 | 0.73 | 11 | 47 | 10 | 0.83 |
| TSA | NP | 22 | 109 | 22 | 0.80 | 6 | 30 | 5 | 0.77 | 5 | 33 | 6 | 0.74 | 9 | 49 | 10 | 0.79 |
|  | LP | 21 | 110 | 22 | 0.79 | 5 | 29 | 7 | 0.80 | 4 | 35 | 5 | 0.65 | 13 | 46 | 9 | 0.85 |
| TRV | NP | 17 | 114 | 22 | 0.74 | 5 | 31 | 5 | 0.73 | 4 | 35 | 5 | 0.65 | 9 | 48 | 11 | 0.81 |
|  | LP | 21 | 107 | 25 | 0.82 | 6 | 27 | 8 | 0.87 | 3 | 35 | 6 | 0.64 | 11 | 48 | 9 | 0.81 |
| RAD | NP | 17 | 113 | 23 | 0.75 | 9 | 25 | 7 | 0.94 | 4 | 35 | 5 | 0.65 | 12 | 45 | 11 | 0.87 |
|  | LP | 28 | 101 | 24 | 0.88 | 5 | 29 | 7 | 0.80 | 6 | 29 | 9 | 0.87 | 10 | 52 | 6 | 0.70 |
| TRT | NP | 18 | 118 | 17 | 0.69 | 3 | 33 | 5 | 0.62 | 7 | 29 | 8 | 0.88 | 8 | 51 | 9 | 0.73 |
|  | LP | 25 | 105 | 23 | 0.84 | 8 | 25 | 8 | 0.94 | 6 | 34 | 4 | 0.69 | 11 | 47 | 10 | 0.83 |
| RF | NP | 18 | 110 | 25 | 0.78 | 5 | 27 | 9 | 0.86 | 7 | 32 | 5 | 0.77 | 6 | 51 | 11 | 0.72 |
|  | LP | 22 | 109 | 22 | 0.79 | 5 | 29 | 7 | 0.80 | 8 | 29 | 7 | 0.88 | 9 | 50 | 9 | 0.76 |

NP: Normal phosphorus; LP: Low phosphorus; PRL: Primary root length; TRL: Total root length; TSA: Total root surface area; TRV: Total root volume; RAD: Root average diameter; TRT: Total root tips; RF: Root forks

The Shannon-Weaver diversity index (H’) was calculated to study the diversity among the tested traits in different genotypic groups (Table 5). The H’ values varied for traits PRL, TRL, TSA, TRV, RAD, TRT and RF with an average of 0.800 studied mungbean genotypes. Among the studied groups it was maximum for Indian Released Varieties. Under NP condition, PRL and TSA exhibited higher H’ value in studied mungbean genotypes. Whereas in LP condition TRL, TRV, RAD, TRT and RF revealed higher H’ value. Under both the phosphorus regimes, RAD and PRL showed relatively higher level of variation while TRV and RF were less variable across different genotypic groups. All root traits except PRL and TSA showed higher H’ values indicate higher diversity under the LP condition than the NP condition. For three traits, TRL, RAD and RF under NP condition and three traits, TRL, TRV and TRT under LP condition showed higher diversity in the IRV group than ABL and G group.

### Principal component analysis

Principal component analysis was carried out to know the most contributing traits under two phosphorus regimes. The first two principal components (PCs) explained the 79.19 % and 78.84 % of the total variation among the tested mungbean lines under NP and LP conditions (**Table 6** and **Fig. 3**). The first principal component explained the 61% and 59% of total variation under NP and LP condition, revealed that TRL and TSA, and their highly correlated traits TRV and RF are the most important contributing traits. The most important contributing trait in second principal component is RAD, which contributed nearly 20% of total variation.

**Table 6:**
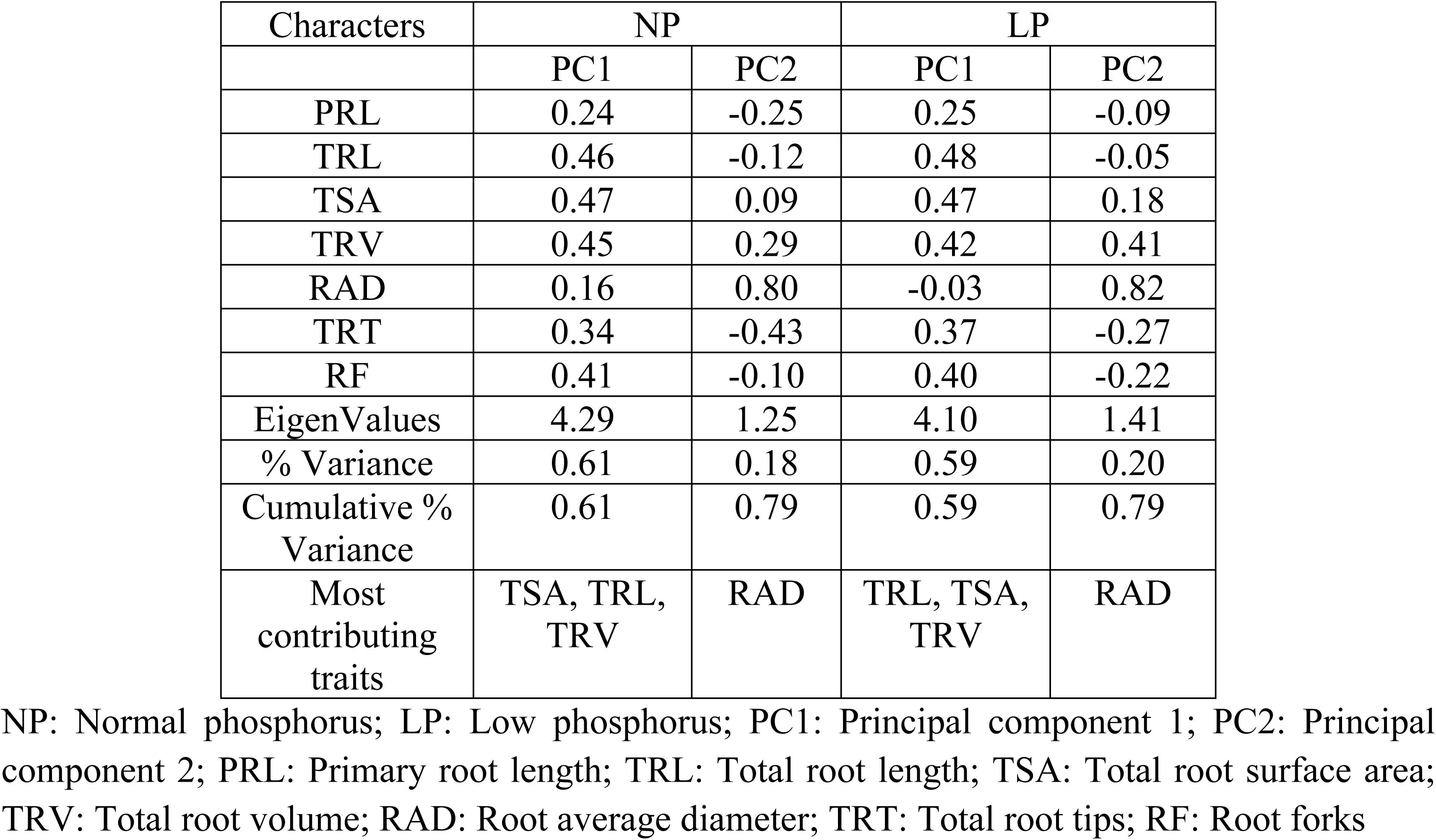
Principle component analysis of seven traits under two phosphorus conditions.

| Characters | NP |  | LP |  |
| --- | --- | --- | --- | --- |
|  | PC1 | PC2 | PC1 | PC2 |
| PRL | 0.24 | -0.25 | 0.25 | -0.09 |
| TRL | 0.46 | -0.12 | 0.48 | -0.05 |
| TSA | 0.47 | 0.09 | 0.47 | 0.18 |
| TRV | 0.45 | 0.29 | 0.42 | 0.41 |
| RAD | 0.16 | 0.80 | -0.03 | 0.82 |
| TRT | 0.34 | -0.43 | 0.37 | -0.27 |
| RF | 0.41 | -0.10 | 0.40 | -0.22 |
| EigenValues | 4.29 | 1.25 | 4.10 | 1.41 |
| % Variance | 0.61 | 0.18 | 0.59 | 0.20 |
| Cumulative % Variance | 0.61 | 0.79 | 0.59 | 0.79 |
| Most contributing traits | TSA, TRL, TRV | RAD | TRL, TSA, TRV | RAD |

NP: Normal phosphorus; LP: Low phosphorus; PC1: Principal component 1; PC2: Principal component 2; PRL: Primary root length; TRL: Total root length; TSA: Total root surface area; TRV: Total root volume; RAD: Root average diameter; TRT: Total root tips; RF: Root forks

### Comprehensive phosphorus efficiency measurement

The P value, as a comprehensive synthetic index was used to study the efficiency of root morphology among mungbean lines under phosphorus deficiency (**Supplementary Table 2**). Based on P values all mungbean lines were classified in to five groups. Group 1 with 21 lines showed highly efficiency for phosphorus uptake with P values greater than 0.9. Group 2 with 25 lines showed efficiency for uptake of phosphorus with P values between 0.7 and 0.9. Group 3 with 48 lines showed moderate efficiency with P values between 0.5 and 0.7. Group 4 with 41 lines showed inefficiency with P values between 0.3 and 0.5. Group 5 with 18 lines showed highly inefficiency with P values less than 0.3. The line which comes under respective groups are listed in **Supplementary Table 2**. The mean values of the phosphorus efficiency coefficient (PC) for each trait in five groups with different levels of phosphorus efficiency are shown in **Fig.4**. The mean values of phosphorus efficiency coefficient (PC) for all root traits were highest in group 1, moderate in group 2, 3 and 4, and lowest in group 5 except for root average diameter (RAD). This result indicates that phosphorus efficient mungbean lines with higher P values also had higher phosphorus efficiency coefficients.

**Fig.3.**
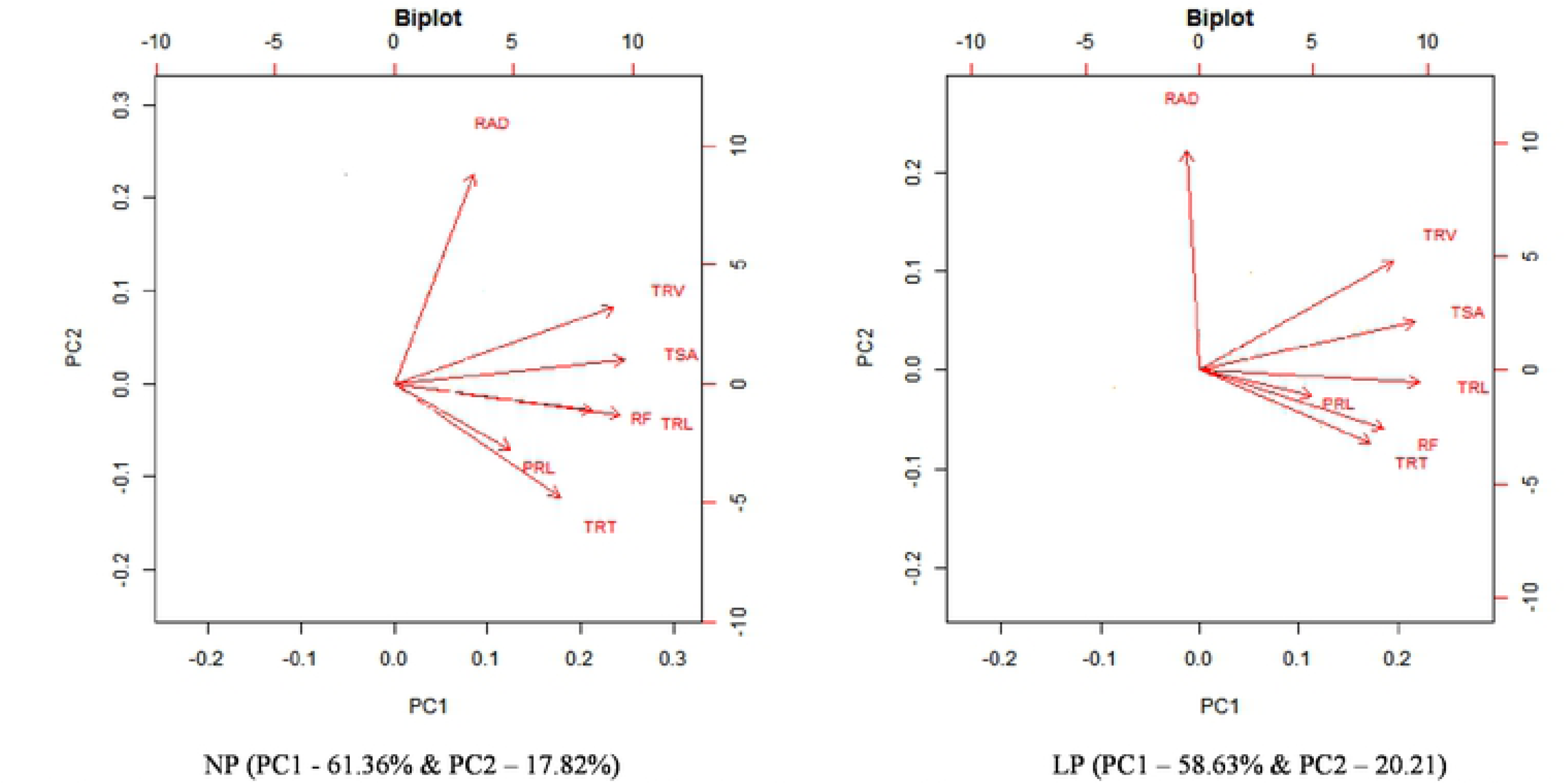
Principal component analysis of seven traits under NP and LP condition. Primary root length; TRL: Total root length, TSA: Total root surface area; RAD: Root average diameter; TRY: Total root volume; TRT: Total root tips; RF: Root forks; NP: Normal phosphorus condition; LP: Low phosphorus condition.

**Fig.4.**
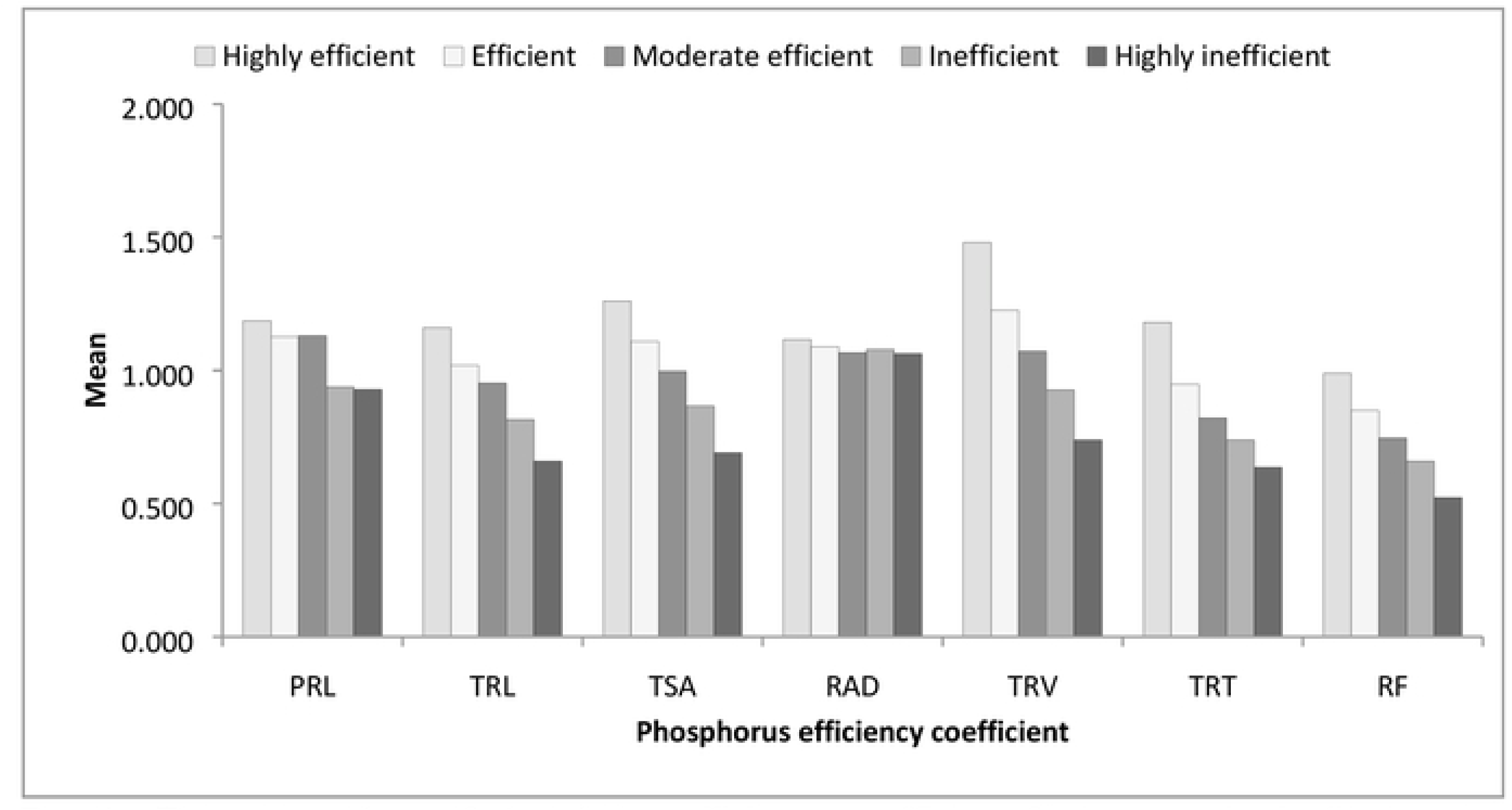
The mean values of phosphorus efficiency coefficient for seven traits in five groups classified for phosphorus efficiency. Groups 1, 2, 3, 4 and 5 represent mungbean lines identified with highly efficiency, efficiency, moderate efficiency, inefficiency and highly inefficiency. N = 21, 25, 48, 41 and 18 for groups 1, 2, 3, 4 and 5 respectively. PRL: Primary root length; TRL: Total root length, TSA: Total root surface area; RAD: Root average diameter; TRY: Total root volume; TRT: Total root tips; RF: Root fork.

## Discussion

Roots are versatile structures absorbing water and nutrients from the soil, providing anchorage to the ground and sometimes storing food and nutrients. Both genetic and environmental factors influence the shape and structure of roots. Understanding the root architecture at different levels of phosphorus is important to our future food security by helping us to breed more phosphorus use efficient varieties. Roots embedded in the soil, are the first organs to respond in low-resource system, particularly in low nutrient and water stress areas [41]. At seedling stage, genotypes can be screened for desirable root traits, which are predictor tools for later stage performance [42]. Root system growth and rhizosphere have a significant role to play for nutrient absorption, mobilization and efficient use by plants [43].

In plants, phosphorus is essential for many metabolic processes including energy transfer, sugar metabolism, intracellular signal transduction, macromolecular biosynthesis, respiration and photosynthesis [44]. Phosphorus deficiency mainly results in poor seedling emergence, slow seedling growth, chlorosis and mature stunted plants at early stages [45] and affects the seed development and fruit maturity at later stages in the growing season [46]. In the present study, veracity of root system was maintained by growing the tested mungbean lines in hydroponic culture. The *in vitro* screening method of hydroponics proves to be ideal method to screen large set of genotypes with least affect of environmental influence [47]. Further, hydroponic system permits the fast screening of root traits of young seedling for traits like nutrient use efficiency [48]. Previous reports examined the root traits using simple hydroponic system based on aerated nutrient solution with different levels of phosphate with replacement of solution at fixed interval in maize [49], soybean [50], mungbean [23], and wheat [51]. Thus root system response to phosphorus deficiency can be studied without root damage in hydroponic culture by controlling access to water and nutrients.

The rate of nutrient uptake by plant roots depends upon the particular nutrient concentration at root surface, root properties and plant requirements [52]. Enhanced root morphological traits, root growth and greater root biomass of phosphorus efficient mungbean lines are connected to the search for particular nutrients under stress conditions. Under low phosphorus conditions, plants modify their root architectural traits [53, 20] which includes reduced primary root growth, increase in number and length of lateral roots and root hairs [54-56], increase in root surface area and volume [24], shallower root growth angle [57] and enhancement of root biomass [58] are the key modifications for enhancement of phosphorus uptake. Characterization of mungbean germplasm lines for stress tolerance traits and screening for phosphorus uptake efficient lines are indispensable for success of breeding programme.

Conventionally, higher root to shoot ratio has been considered as index for phosphorus efficiency due to increase in root biomass and large deep root system able to extract more nutrients [59, 60]. Total root length represents the sum of primary, seminal, crown, basal and lateral roots. The various components of root system have also been selected as important traits for screening of lines under phosphorus deficiency. In this study, we examined the influence of phosphorus deficiency on root morphology of 153 mungbean lines and investigated the various root traits including PRL, TRL, TSA, TRV, TRT and RF. We found the significant variation, medium to high heritability, approximately normal distribution and significant correlations for these root traits. Although, genetic variation of root system vary from plant to plant, the presence of very fine roots (<0.5 mm diameter) and fine roots (0.5 to 2.0 mm) determines the most percentage of root traits is important for nutrient and water uptake [61-63]. In this study, we identified the high percentage of fine roots with diameter from 0.5 to 2.0 mm in LP compared to NP conditions, while percentage of very fine roots with <0.5 mm diameter more in NP condition. This indicates the affect of phosphorus availability on percentage of fine root distribution at different diameter classes in studied mungbean genotypes. Under low phosphorus conditions, plant may increase the development of root cortical aerenchyma which enables the plant to maintain greater root diameter but reduce overall total root cost and root respiration [64, 65].

PCA analysis showed that TRL, TSA, TRV and RAD were responsible for most of the phenotypic variation at seedling stage in the tested mungbean lines. TRL was significantly and positively correlated with TSA, TRV, TRT and RF under both NP and LP conditions. In combination with PCA analysis, we identified that TRL, TSA and TRV were sufficient to explain the most of variation and these were proved to be ideal traits for phosphorus uptake efficiency screening at seedling stage. Under LP condition, root average diameter was significantly and negatively correlated with total root tips and root forks. This indicates that root average diameter is a key trait to differentiate phosphorus availability among the tested root traits. Moreover, these traits showed high phosphorus efficiency coefficient values in phosphorus efficient mungbean lines. This result is in agreement with previous reports. Pandey *et al*. [23] reported the significant higher root surface area and root volume in phosphorus efficient mungbean genotype under phosphorus stress. Root surface area has been found to be close association with nutrient absorption rate [66, 67]. Vigorous root growth with high root length and surface area ensures the efficient absorption of macro and micronutrients at early growth stage of plant [68]. Furthermore, root architectural traits mainly total root length and root number were significantly and positively correlated with biomass and grain yield [69, 70]. Therefore, vigorous root system of plant not only supports good crop establishment but also ensure the plant survival under stressful conditions.

Diversity in root architecture enables us to improve nutrient and water use efficiency under stressful conditions. A combination of availability of diverse mungbean genotypic lines and stress tolerance ability will be key criteria for success of crop improvement programme. In this study, comparison of root morphological traits across different genotypic groups indicated that IRV group showed greater diversity for root traits than ABL and GL groups. Furthermore, Shannon–Weaver diversity index (H’) was calculated to compare the phenotypic diversity among the traits. Among all traits, TRL and RAD showed relatively high level of H’ under LP condition and PRL showed relatively highest value of H’ under both phosphorus regimes. High value of H’ indicates greater genetic diversity and balanced frequency distribution [71], while low H’ indicates extreme unbalanced frequency distribution with lack of diversity [72]. This result provides valuable information to improve both agronomic traits as well as nutrient use efficiency traits in mungbean breeding programme. In the 21^st^ century, due to environmental concerns and high cost of inorganic fertilizers, nutrient efficient crop plants play an important role in improving crop yields compared to 20^th^ century [73].

Based on comprehensive index of P values, 21, 25, 48, 41 and 18 mungbean lines were classified as highly efficient, efficient, moderately efficient, inefficient and highly inefficient groups respectively. Among these, IPM-288, TM 96-25, TM 96-2, M 1477, PUSA 1342 were identified as best five highly efficient genotypes whereas M 1131, PS-16, Pusa Vishal, M 831, IC 325828 were highly inefficient genotypes. Except RAD, phosphorus efficiency coefficients for all traits were highest in group 1, intermediate in 2, 3 and 4 and lowest in group 5. This type of classification is required for screening and selection of genotypes for desirable root traits under varied phosphorus conditions. Further, these genotypes with contrasting traits can be exploited in recombination breeding programme to develop phosphorus efficient cultivars [74, 75]. In this study, 21 high efficient lines with well developed root system were identified and these could be used in mungbean breeding programme for further improvement of tolerance to abiotic stresses.

In conclusion, we identified a range of response to phosphorus deficiency in mungbean lines for root system traits at the seedling stage. We found that TRL, TSA and TRV are the ideal selection criteria at seedling stage for predicting the nutrient use efficiency in the field. Further, the tested mungbean lines needs to be evaluated at the adult stage under NP and LP conditions. In addition, association of seedling stage root traits with adult stage traits needs to be further examined.

